# Solubility-aware protein binding peptide design using AlphaFold

**DOI:** 10.1101/2022.05.14.491955

**Authors:** Takatsugu Kosugi, Masahito Ohue

**Affiliations:** Department of Computer Science, School of Computing, Tokyo Institute of Technology, G3-56-4259 Nagatsutacho, Midori-ku, Yokohama City 226-8501, Kanagawa, Japan

**Keywords:** Solubility, Peptide design, AlphaFold, AfDesign

## Abstract

New protein–protein interactions (PPIs) are being identified, but PPIs have different physicochemical properties compared with conventional targets, making it difficult to use small molecules. Peptides offer a new modality to target PPIs, but designing appropriate peptide sequences by computation is challenging. Recently, AlphaFold and RosettaFold have made it possible to predict protein structures from amino acid sequences with ultra-high accuracy, enabling de novo protein design. We designed peptides likely to have PPI as the target protein using the “binder hallucination” protocol of AfDesign, a de novo protein design method using AlphaFold. However, the solubility of the peptides tended to be low. Therefore, we designed a solubility loss function using solubility indices for amino acids and developed a solubility-aware AfDesign binder hallucination protocol. The peptide solubility in sequences designed using the new protocol increased with the weight of the solubility loss function; moreover, they captured the characteristics of the solubility indices. Moreover, the new protocol sequences tended to have higher affinity than random or single residue substitution sequences when evaluated by docking binding affinity. Our approach shows that it is possible to design peptide sequences that can bind to the interface of PPI while controlling solubility.

## 1. Introduction

Protein–protein interactions (PPIs) are associated with various biological processes, such as signal transduction and metabolism, and play a fundamental role in many cellular activities [1]. Approximately 810,000 human non-redundant PPIs have been identified [2] (BioGrid 4.4.208 - April 2022). These PPIs regulate many important intracellular pathways in human diseases and have attracted attention as potential drug targets since the early 2000s [3–7]. There have been several successful small-molecule drug designs targeting PPIs, including the Federal Drug Administration (FDA)-approved PPI inhibitor Venetoclax [8], which is considered a BH3 mimetic, and others in the clinical stage [9,10].

However, the physicochemical properties of PPI drug targets are significantly different from those of conventional drug targets, and PPI-targeting drug discovery remains a severe challenge [9,11]. One of the most important reasons for the difficulty in PPI-targeting drug development is shallow and large PPI interfaces. Small molecule binding sites are relatively deep and have an area of only approximately 500 Å2, while PPI binding interfaces are typically flat and wide, with a surface area of approximately 1,000–4,000 Å2 [12–14]. Another difficulty in PPI-targeting drug discovery is that small molecule drugs tend to bind to the deep folding pockets of proteins rather than the large, flat, and, hydrophobic binding interface of the PPI [15]. Antibodies are an effective modality for recognizing PPI interfaces; although they are effective for targeting receptors on the cell membrane surface, they are not effective against intracellular PPI targets because they have difficulty penetrating the cell membrane [16].

In recent years, peptides with a balance of flexibility and binding affinity have attracted attention as a modality for targeting PPIs. Research on therapeutic peptides began with natural human hormones such as insulin, vasopressin, oxytocin, and gonadotropin-releasing hormone. To date, approximately 80 peptide drugs have been approved, with approximately 30 non-insulinic peptide drugs approved since 2000. More than 170 peptides have been in clinical development. As of 2021, there are 58 therapeutic peptides targeting PPIs in clinical stages, of which 13 are in Phase 1, 26 in Phase 2, 15 in Phase 3, and 4 have New Drug Applications pending and are close to approval [17,18].

There are several methods for experimental design of PPI-targeting peptides, such as phage display, mRNA display, and other methods for screening large numbers of peptides that interact with proteins experimentally, but the cost is very high [19–21]. Therefore, a method is needed to computationally design candidate peptide sequences that interact with target proteins. While there are examples of designing sequences that have a function, such as the design of antimicrobial peptide sequences [22,23], a lot of experimental data is needed, and it is difficult to obtain a lot of data on peptide sequences that interact with specific proteins targeting PPIs. There is also a method called PDGA that uses genetic algorithms to generate peptide sequences with similar small molecule footprints, but this method also requires at least one example of a small molecule compound that is the target of a specific protein, and the footprints do not consider actual 3D structure [24]. In general, protein designs are designed using the three-dimensional (3D) structure of a protein that binds to a specific site on the surface of the target protein [25–28]. For more efficient design of PPI-targeted peptides, it is important to utilize conformational information on the target protein and the PPI-binding interface.

Recently, protein structure prediction using deep neural networks, which estimate 3D structures from the co-evolution of amino acid residues, has been widely used [29,30]. It is also said that AlphaFold can be used for modeling peptide structures smaller than 60 amino acids if the target peptide has a well-defined secondary structure and does not have multiple turned regions or flexible regions that may take different conformations [31]. Furthermore, it has been shown that docking of proteins and peptides is possible using AlphaFold [31]. Furthermore, it has recently become possible to predict sequences from protein structures. For example, it was reported that Monte Carlo sampling in amino acid sequence space was used to design sequences with specific backbone structures from random sequences. Moreover, a pipeline has been developed to design small proteins that bind to target proteins based on the results of many experiments [33,34]. Similarly, methods have been published to predict the backbone of a specific 3D structure or the sequence of a complex structure of a target protein from a random sequence using all-atom geometry and AlphaFold prediction accuracy indicators, such as the predicted aligned error (PAE) and predicted local-distance difference test (pLDDT) [35,36]. Protein design has become possible using such deep learning networks.

AfDesign implements the “binder hallucination” method, which is based on the AlphaFold peptide docking method and the new protein design method “deep network hallucination” described above. However, our examination of peptide design using AfDesign binder hallucination showed that the sequences to be designed tend to have more aromatic ring and hydrophobic residues on the interaction surface, and the logS, an index of water solubility of the designed peptide sequences, tends to be relatively low.

The development of strategies to improve membrane permeability and facilitate cellular uptake will be essential for successful targeting of PPIs, because peptides have low cell membrane permeability [38]. One way to improve bioavailability is through water solubility. The higher the water solubility, the easier it is to maintain effective serum concentrations, and thus bioavailability is usually greatly enhanced. High solubility is also one of the requirements for successful recombinant protein production and is an important industrial factor [39]. Although water solubility is an optimization issue in the design of therapeutic peptides, experimentally identifying unnecessary hydrophobic amino acids and replacing them with charged or polar residues adjusts the isoelectric point while maintaining biological activity, which is said to be primarily an empirical process [40,41]. Recently, there have been many attempts to computationally solve the empirical process by using machine learning to predict protein solubility [42].

There are several possible strategies for peptide design to target PPIs considering solubility.

- A library of highly water-soluble peptide sequences is created, and then docking scores and binding affinities are predicted by protein–peptide docking.
- Design peptide sequences that are likely to bind to target proteins using peptide sequence prediction methods such as AfDesign, then evaluate water solubility and filter out those that exceed water solubility thresholds.

The first strategy is that the number of sequences in the amino acid sequence of *N* residues has a huge pattern of 20*N*, and even if we set some constraints based on the characteristics of the amino acid residues in some way, it is computationally expensive and impractical to search for the target peptide from the large number of candidate peptides by docking calculations. The second strategy is that, as mentioned above, owing to the characteristics of AfDesign, binder hallucination tends to increase the number of aromatic ring residues and hydrophobic residues on the interaction surface, and therefore, after design, there is a possibility that few candidate sequences will remain when filtered using water solubility indices such as logS. The reason why the above strategy does not work well may be because peptide design and solubility optimization are not performed at the same time.

Therefore, this research focused on the possibility of controlling solubility while maintaining or improving the binding affinity of the designed peptide by simultaneously optimizing the binder peptide sequence of the target protein and optimizing solubility by designing the generation process of AfDesign with a loss of solubility constraint.

## 2. Materials and Methods

### 2.1. AfDesign settings

We used the AfDesign binder hallucination protocol to calculate the binder of the MDM2 protein. The Protein Data Bank (PDB) 1YCR chain A was used as the MDM2 structural template for AfDesign. The binder_len is the model of the p53 peptide in 1YCR. The length was set to 13, the same as the length of the sequence “ETFSDLWKLLPEN”. The length of the sequence was set to 17 when AfDesign was applied with PD-1 as the target protein. The design method used was design_3stage(); soft_iter, temp_iter, and hard_iter were set to 100, 100, and 10, respectively. Unless otherwise noted, other settings were left at default. AfDesign was run 100 times for all three solubility indices with 100 different settings of seed from 1 to 100 for each weight (0.1, 0.3, 0.5, 0.7, 1.0) when added to the other weights in AfDesign. The same was done for the conditions without a solubility index. The versions of jax and jaxlib used in this study were 0.3.1 and 0.3.0+cuda11.cudnn805, respectively. To ensure reproducibility, TF_CUDNN_DETERMINISTIC=1 was set JAX to behave deterministically. While AfDesign was updated during our study, the commit history we used was that of: ColabDesign repository: https://github.com/sokrypton/ColabDesign/tree/be242491a2517589c96f09b7d15b7bb37f72bafa af_backprop repository: https://github.com/sokrypton/af_backprop/tree/7246fe544e9398de8dab848b11b7b634f16858db The AlphaFold model used in AfDesign was downloaded from https://storage.googleapis.com/alphafold/alphafold_params_2021-07-14.tar

### 2.2. Solubility loss calculation

Three solubility indices, the Hydrophobicity Index [43], Hydropathy Index [44], and Solubility-Weighted Index (SWI) [42], were used in this study (Supplementary Table S1). The Hydrophobicity Index evaluates hydrophobicity based on the physical characteristics of 20 amino acids to identify regions of a protein’s primary sequence that are likely to be buried in the membrane [43]. The Hydropathy Index is a hydrophilicity scale that considers the hydrophilicity and hydrophobicity of each of the 20 amino acid side chains and was developed based on experimental observations from the literature. Specifically, values were calculated using both the water vapor transfer free energy and the distribution in and out of the amino acid side chains as determined by Chothia (1976) [44]. The Solubility-Weighted Index is a predictive index of solubility, and prediction programs using it are superior to many existing de novo protein solubility prediction tools [42]. In this study, the weight of this predictive index was used as the solubility index.

The three solubility indices were normalized to have a maximum of 1 and a minimum of 0. Owing to the meanings of the indices, the normalized Hydrophobicity and Hydropathy indices were used as they are in the solubility loss function. For the normalized SWI, values inverted (minus 1) after normalization were used for the solubility loss function. All modified index values are shown in Supplementary Table S2.

The solubility loss is the geometric mean of the designed amino acids and the corresponding normalized solubility index values. A solubility loss was added to the other losses specified in AfDesign.

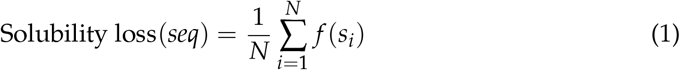

where *seq* represents the designed amino acid sequence and *seq* = (*s*_1_*s*_2_…*s*_*N*_), *s*_*i*_ represents an amino acid residue in the *i*-th position of *seq, N* is the sequence length of the binder_len to be designed, *f* represents the amino acid index, and *f* (*s*) is the index value of the amino acid residue *s*.

### 2.3. Calculation of solubility

As a preprocessing step, SMILES were prepared by acetylation of the N-terminus of the peptide sequence and amidation of the C-terminus using RDKits Chem.MolFromHELM(), and conformers were prepared by LigPrep in the Schrödinger software suite (version 2021-1). Finally, logS values were calculated as solubility using QikProp in the Schrödinger software suite (version 2021-1).

### 2.4. Calculation of protein–peptide binding affinity

Using AutoDock CrankPep within AutoDockFR software suite version 1.0 [45], 1YCR chain A as a receptor, 1YCR chain B, and the designed peptide as a peptide were calculated. The procedure is as described in the official document. First, the PDB files prepared as receptor and peptide were protonated by reduce version 3.23.130521 [46], and then the protonated PDB files were converted to PDBQT files by using two scripts, prepare_ligand and prepare_receptor, of the AutoDockFR software suite. In addition, a docking box was placed and sized over the receptor pocket into which the peptide was to be docked and a trg file was calculated using agfr to compute an affinity map for the list of atom types in AutoDock 4 [47]. Up to 2.5 million evaluations of the scoring function were performed to increase the likelihood of finding the best docking pose (the global minimum of the scoring function) and 50 independent searches were performed for each. The argument of AutoDock CrankPep is -N 50 -n 2500000.

### 2.5. Creation of sequence logos

The sequence with the lowest final loss in AfDesign in the latter 10 iterations (hard_iter) design_3stage() was selected as the best sequence. Sequence logos were created on the WebLogo 3 server (http://weblogo.threeplusone.com/create.cgi) [48] with the 100 best sequences per solubility index and per weight as input.

## 3. Results

### 3.1. Binder design targeting PPI using AfDesign

Our approach to solubility-aware protein-binding peptide design was performed using the AfDesign binder hallucination. AfDesign is a public module created by Dr. Sergey Ovchinnikov [36], and is a tool for designing protein-binding peptides based on Dr. Sergey’s previous work with trRosetta [33,49]. AlphaFold is used as the structure prediction oracle, the optimization target is the binding sequence, and the loss function is defined as a flexible function of AlphaFold confidences. It returns the pLDDT, PAE, and distogram, among which pLDDT measures the quality of the local model for each residue, PAE indicates the confidence level of each amino acid residue pair, and distogram indicates the distance prediction probability of the C_*β*_ of an amino acid residue pair. The objective of this study was to investigate whether a new loss function with a solubility measure could be added to the original AfDesign loss function to optimize the solubility of the sequence to be designed. Binding affinity was measured using AutoDock CrankPep and further validated by measuring logS by QikProp as a measure of solubility. A schematic diagram of the whole process is shown in Figure 1.

**Figure 1.**
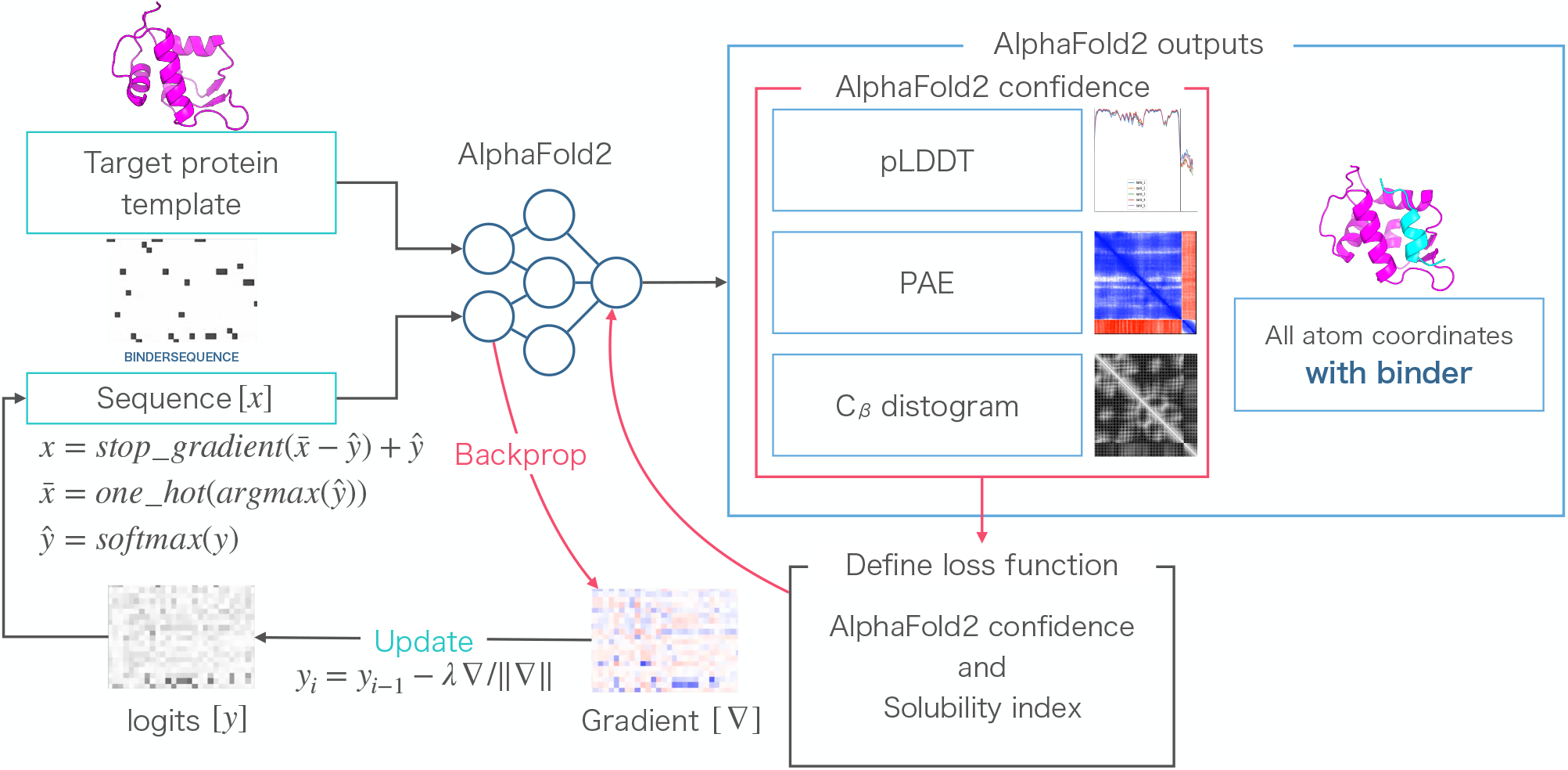
Schematic of the optimization method of the AfDesign binder hallucination. The output of AlphaFold is an all atom coordinate, two reliability indices (predicted aligned error [PAE] and predicted local-distance difference test [pLDDT]), and a distogram. These are used to define the loss function, which is back-propagated to compute the gradient for the design sequence, and then updated and predicted in a loop for optimization. To evaluate the optimized sequence, the binding affinity is calculated by docking and logS is calculated as a peptide solubility index.

The principle of the AfDesign binder design is that the PDB file of the target protein is used as a template, and the PDB file is matched to the AlphaFold input. In addition, the “residue index” of the last residue of the target protein and the starting residue of the complex peptide are sufficiently spaced to create a “chain break” state [50]. This is a trick to predict complexes using AlphaFold and is also used in ColabFold [51]. The “chain break” peptide is a random sequence with a specified length and an initial sequence. Finally, a loss function is created using the confidence index output from the AlphaFold model, which is then used as a backprop to optimize the “chain break” peptide sequence as a complex [36]. We used the MDM2-p53 complex (PDB ID: 1YCR) in this research as representative example of a PPI protein complex. The target protein was MDM2, and the original binder was p53 peptide.

To evaluate the distribution of solubility of the binder sequences designed in the AfDesign binder hallucination protocol, 100 designs were performed using MDM2 as the target protein and different seed settings in AfDesign. The logS of the p53 peptide sequence, a known binding peptide of MDM2, and the binder sequence designed in the AfDesign binder hallucination protocol were calculated by Schrödinger’s QikProp as an index of solubility. As shown in Figure 2(a), the distribution of logS of the binder sequence designed in the AfDesign binder hallucination protocol was much lower than that of the p53 peptide sequence, indicating that the binder tends to be designed with lower solubility. To confirm the amino acid composition of the designed sequence, the sequence logos were generated from 100 designed sequences using WebLogo [48]. Figure 2(b) shows that the binder sequences designed in the AfDesign binder hallucination protocol contained more hydrophobic amino acid leucine and aromatic amino acids tyrosine, phenylalanine, and tryptophan in residues 4 to 11, In general, the binding interface of PPI was consistent with high hydrophobicity [52]. Comparing the designed sequence with the sequence of the p53 peptide, “ETFSDLWKLLPEN”, the composition ratio of hydrophobic amino acids tended to be higher and the solubility lower.

**Figure 2.**
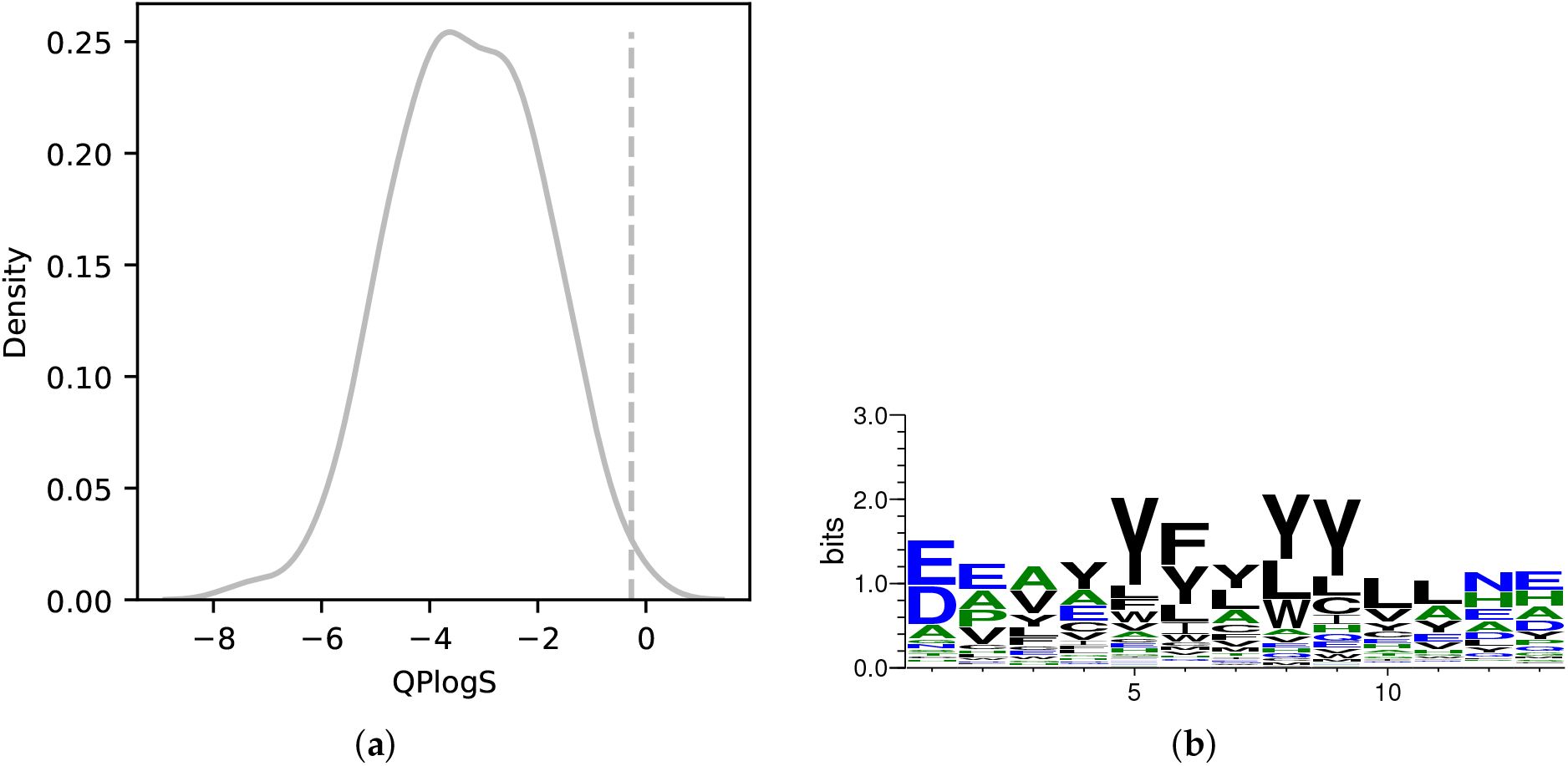
Distribution of logS of MDM2 binder sequences designed by AfDesign and sequence logo of the binder sequences. Distribution of logS calculated by QikProp. Dashed lines are logS values for the p53 peptide (**a**). Sequence logo for sequences designed with the AfDesign binder hallucination protocol (**b**). Hydrophilic amino acids (R, K, D, E, N, Q) are shown in blue, neutral amino acids (S, G, H, T, A, P) are shown in green, and hydrophobic amino acids (Y, V, M, C, L, F, I, W) are shown in black.

From the results in Figure 2(b), the logS distribution of the binder sequence designed with the default AfDesign binder hallucination protocol was relatively low compared with that of the p53 peptide. Therefore, in order to test whether it is possible to control water solubility from the bioavailability and industrial perspective of therapeutic peptides by adding solubility constraint as a new loss to the design process of the AfDesign binder sequence, we simultaneously optimized the binder peptide sequence of the target protein, MDM2, and optimized the solubility. Based on previous studies on amino acid and protein solubility, three solubility indices were used to verify the results, the:

- Hydrophobicity Index [43]
- Hydropathy Index [44]
- Solubility-Weighted Index [42]

The different index values related to solubility were each normalized and added as solubility loss to the loss function for the AfDesign binder hallucination protocol, and the weights of the loss were weighted in the range of 0.0–1.0 to design the binders. The following is a summary of the results.

### 3.2. Solubility-aware binder design using AfDesign

As shown in Figure 3, the sequence designed using SWI for solubility loss had the lowest degree of change in solubility with weight among the three indices, while the sequence designed using the Hydrophobicity Index as the solubility index had the highest degree of change in solubility with weight among the three indices. The sequence logo of the sequence designed with the AfDesign binder hallucination protocol considering solubility loss is shown in Figure 4. As Figure 4 shows, compared with the Hydropathy Index, the aromatic amino acids tyrosine and phenylalanine gradually decreased in sequence frequency with weight in the Hydrophobicity Index and SWI. This is consistent with the greater weight of tyrosine and phenylalanine. Moreover, compared with the Hydrophobicity Index and SWI, the Hydropathy Index showed a gradual decrease in sequence frequency of leucine, a hydrophobic amino acid, along with its weight. This is consistent with the greater weight of leucine (see Supplementary Table S1). The addition of solubility loss to the AfDesign binder hallucination protocol resulted in an increase in solubility, while the sequences were designed according to the characteristics of each solubility index.

**Figure 3.**
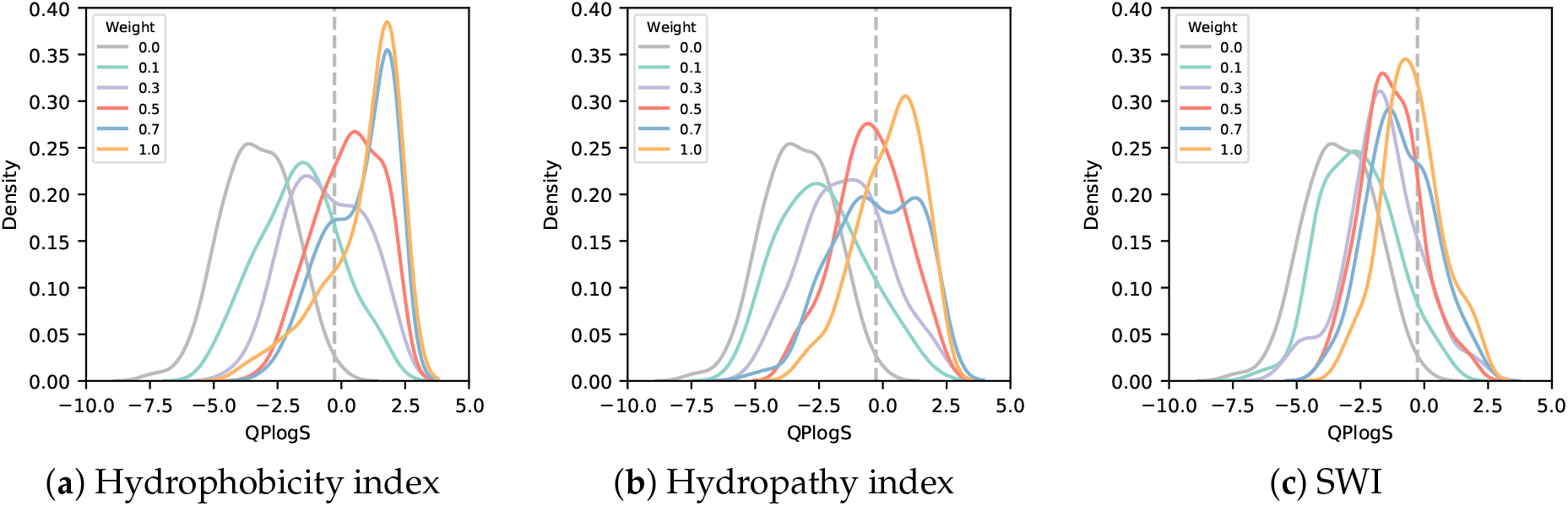
Distribution of logS for various weights of sequences designed in AfDesign using three solubility indices. The logS of sequences designed in AfDesign using the Hydrophobicity Index (**a**), Hydropathy Index (**b**) and Solubility-Weighted Index (**c**) as solubility indices in the loss function. Each color indicates the weight of the solubility loss. The gray line with a weight of 0.0 shows the distribution of logS for the sequence designed by AfDesign without a solubility index. The dashed line shows the logS of the p53 peptide.

**Figure 4.**
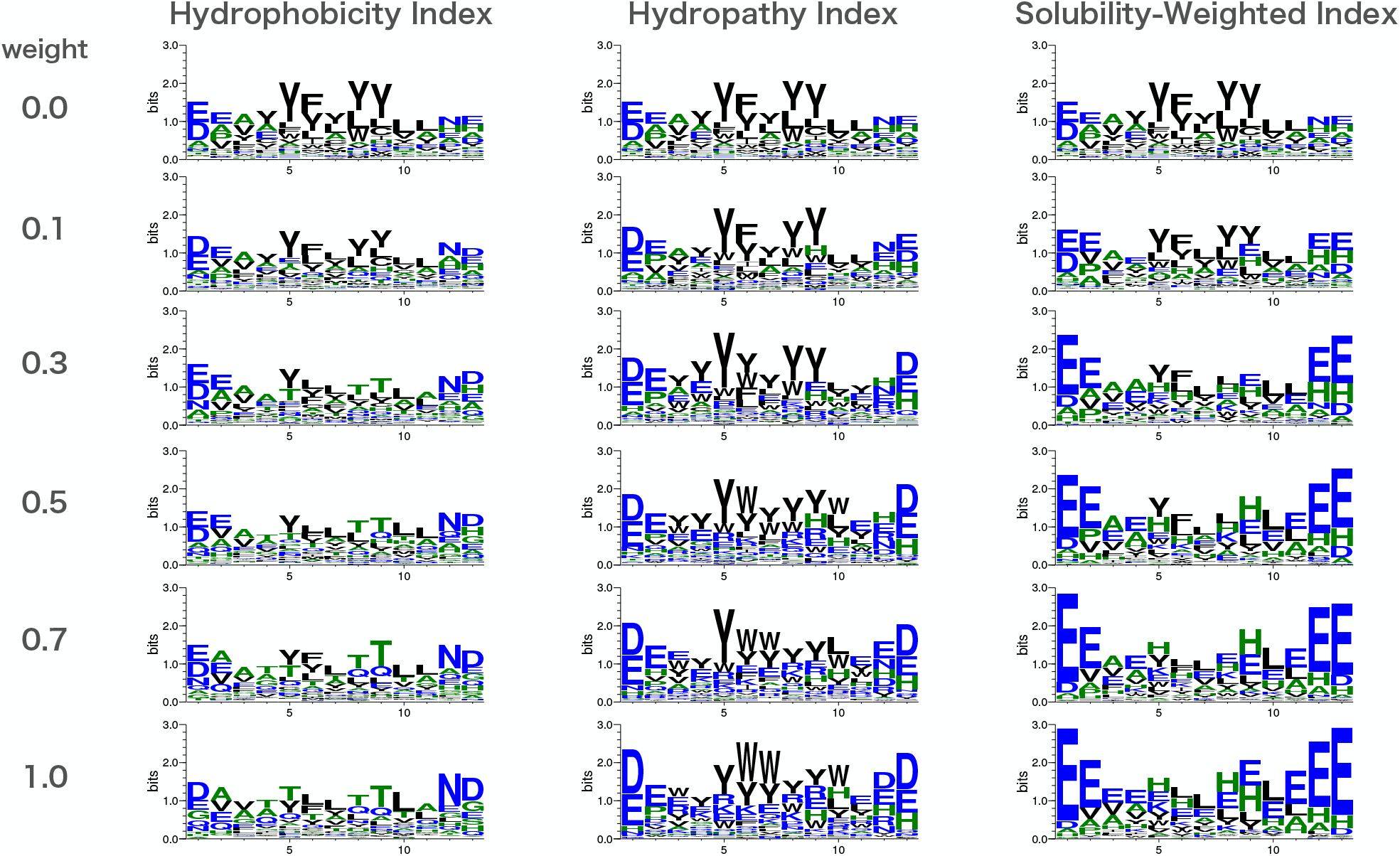
Sequence logos designed by AfDesign binder hallucination with solubility loss for each weight parameter. Hydrophilic amino acids (R, K, D, E, N, Q) are shown in blue, neutral amino acids (S, G, H, T, A, P) are shown in green, and hydrophobic amino acids (Y, V, M, C, L, F, I, W) are shown in black.

To assess the binding affinity of sequences designed with the AfDesign binder hallucination protocol with loss of solubility to the target protein, MDM2, we calculated the binding affinity of the designed peptide to the target protein using AutoDock CrankPep. As shown in Figure 5, the only sequence designed using the Hydrophobicity Index had a lower binding affinity to the target protein than the sequence designed without solubility loss. The binding free energy tended to be higher (weaker as a value of binding). Sequences designed using the Hydropathy Index tended to maintain or increase binding affinity compared with sequences designed without solubility loss.

**Figure 5.**
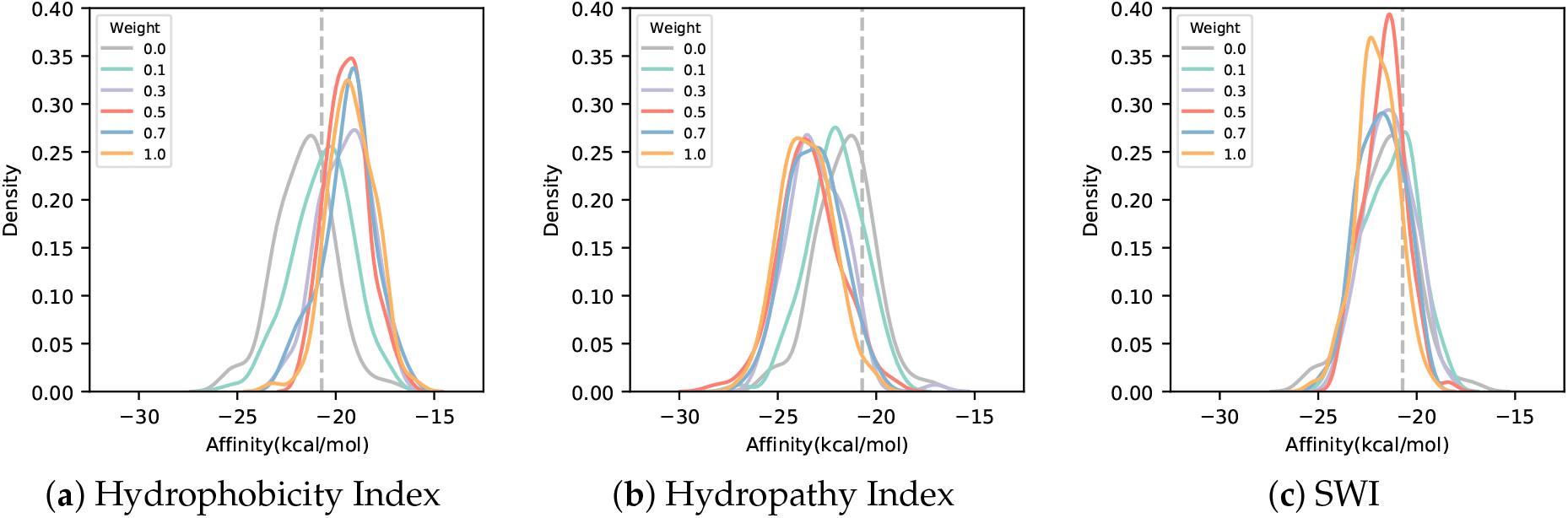
Distribution of binding affinity for various weights of sequences designed in AfDesign using three solubility indices. The binding affinity of sequences designed in AfDesign using the Hydrophobicity Index (**a**), Hydropathy Index (**b**), and Solubility-Weighted Index (SWI) (**c**) as solubility indices in the loss function. Each color indicates the weight of the solubility loss. The gray line with a weight of 0.0 shows the distribution of binding affinity for the sequence designed by AfDesign without a solubility index. The dashed line shows the average of binding affinity of the p53 peptide.

In order to examine the possibility of designing sequences with higher binding affinity and logS than existing complex peptides for the target protein, which was the objective of this study, a scatter plot of binding affinity and logS is shown Figure 6. Figure 6 shows that sequences designed using the Hydrophobicity Index for solubility loss tended to decrease in binding affinity as logS increased, while sequences designed using the Hydropathy Index showed higher binding affinity and higher logS than the sequences designed using the other indices or sequences without solubility loss. The sequences designed using the Hydropathy Index tended to have higher binding affinity with increasing logS than sequences designed using the other indices or sequences without solubility loss. This was the opposite trend for sequences designed using the Hydrophobicity Index. Sequences designed using SWI for solubility loss showed little change in binding affinity as logS increased. Thus, the

**Figure 6.**
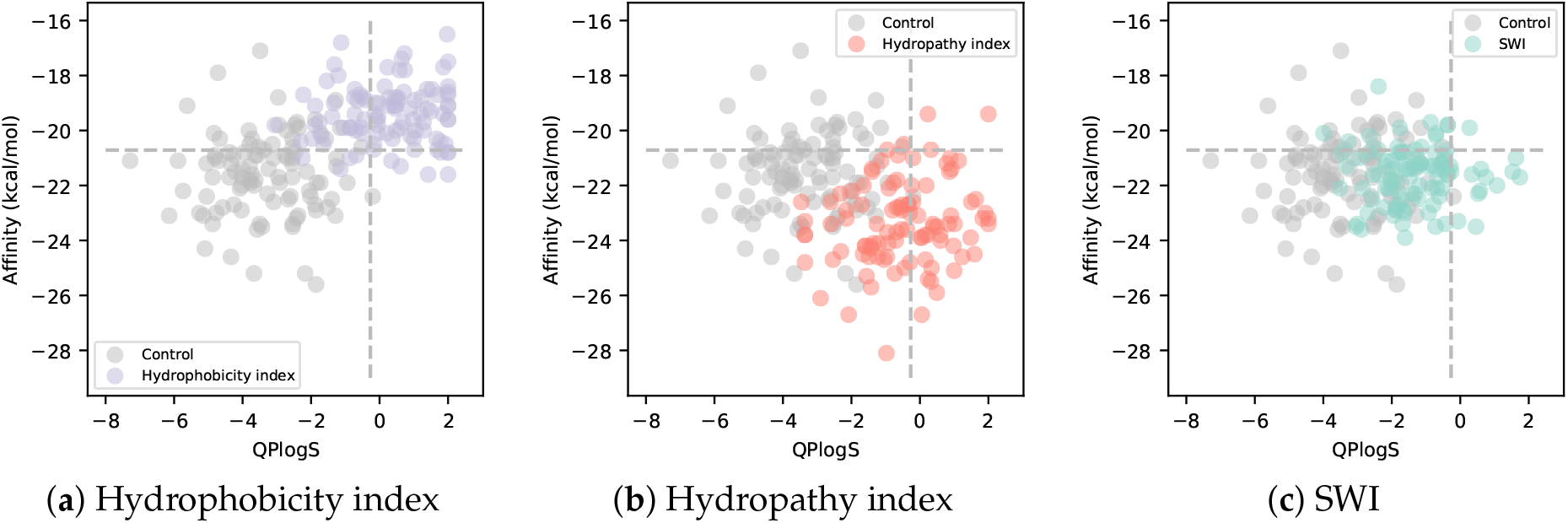
Scatter plots of binding affinity and logS for various weights of sequences designed in AfDesign using three solubility indices. Each point shows the binding affinity and logS of a sequence designed in AfDesign with a weight of 0.5 using the Hydrophobicity Index (**a**), Hydropathy Index (**b**), and Solubility-Weighted Index (SWI) (**c**) as solubility indices in the loss function. Gray dots show the binding affinity and logS of sequences designed in AfDesign without a solubility index. The horizontal dashed line shows the average binding affinity of the p53 peptide. The vertical dashed line shows the logS values for the p53 peptide.

Hydrophobicity Index and SWI were able to control logS while maintaining or increasing the binding affinity of sequences designed by the AfDesign binder hallucination protocol with solubility loss.

## 4. Discussion

The results of this study show that the AfDesign binder hallucination protocol with the addition of the Hydropaty Index or SWI for solubility loss can design candidate peptide sequences that may have binding affinity comparable to or greater than that of the p53 peptide. The binding affinities of sequences designed with the AfDesign binder hallucination protocol were higher than those of random sequences. In addition, the binding affinity of the sequence with one residue substitution of p53 peptide was distributed around the binding affinity value of the p53 peptide; its distribution was lower than that of the binding affinity of the sequence designed with the binder hallucination protocol of AfDesign with or without the Hydropaty Index and SWI for solubility loss (Supplementary Figure S1). In summary, it is likely that sequences designed with the AfDesign binder hallucination protocol have higher binding affinities than random sequences as well as the binding affinity of the p53 peptide and its residue substituted sequences.

In addition to the evaluation of solubility and binding affinity, visualization using PyMOL and evaluation of docking using DockQ were also performed(Supplementary Figure S2). Comparing the 3D structure of the sequence with the highest binding affinity and higher logS than the p53 peptide, which was designed using the solubility loss with SWI in the AfDesign binder hallucination protocol, with the actual 3D structure of MDM2, we find that the coordinates of the sequence are almost identical to those of the p53 peptide.

DockQ was calculated to be 0.500. LRMS, the RMSD of the main chain atoms of the peptide, was calculated to be 5.303; iRMS, the RMSD of the heavy atoms of the residues at the interface between the ligands, was calculated to be 2.294. This is of medium quality in the protein–protein docking criteria of CAPRI and indicates the validity of the sequence designed using solubility loss with SWI in the AfDesign binder hallucination protocol to dock with MDM2.

The impact of adding the solubility index to the loss of the AfDesign binder hallucination protocol is mainly seen in the change in loss from the 101st iteration, when the sequence features are first converted from logits to probabilities by multiplying logits by softmax, and from the 201st iteration, when the probability argmax is first converted to probabilities by multiplying softmax by logits. The main impact on the overall loss of AfDesign is the change in the loss of pLDDT and PAE_inter from the 101st iteration, when the array features are first converted to probabilities by multiplying logits by softmax, and the 201st iteration, when the probabilities are first converted to argmax. This tends to increase the overall loss. Moreover, the variation after the 201st iteration is particularly large compared with the case where the solubility index is not added to the loss of the AfDesign binder hallucination protocol. By varying the solubility loss weights, the solubility loss decreases. This may be a trade-off between optimizing for sequences that tend to have lower solubility according to the solubility index and optimizing for peptide sequences that are likely to form optimal complexes (Supplementary Figure S3).

In terms of designing PPI inhibitor peptides, the question is whether it is possible to design peptides in which the *β*-sheet serves as the PPI binding interface as well as those in which the helix domain serves as the PPI binding interface, as in the MDM2p53 complex. Therefore, we tested the possibility of designing a PD-1PD-L1 complex as an example of a complex in which the *β*-sheets are at the PPI binding interface, and AfDesign is able to design a binder that can form a complex with PD-1. The results show that the default AfDesign binder hallucination protocol is sufficient to design a peptide with high solubility. However, we also found that the distribution of binding affinity was almost the same as that of the random sequence (Supplementary Figure S4).

By creating a solubility loss using three solubility indices for the AfDesign binder hallucination, we were able to design sequences that maintained or increased binding affinity while controlling solubility. The sequences designed by AfDesign could be adversarial, such as those simply preferred by AlphaFold. However, in this study, instead of evaluating the sequences by AfDesign alone, we sampled multiple seeds, calculated the binding affinity to the target protein by the Docking tool, which is not related to AlphaFold, and calculated logS using the solubility calculation tool. The results show an overall trend of the target protein having a high affinity and solubility being controlled. However, these results are based on computer simulations; true solubility and binding affinity will not be known until experiments are conducted. However, the ability to design sequences with high solubility and high binding affinity that could be candidates for PPI inhibition from among a large number of candidate sequences is meaningful from both the drug discovery and industrial perspectives.

Recently, it was reported that competitive binding can be simulated in protein–peptide docking using ColabFold [53]. In other words, it is possible to perform virtual competition binding experiments of peptides. We further performed a computational competition binding test preliminary between the designed peptide with high binding score and the original p53 peptide, and showed that the binding affinity may be higher than that of p53, which is consistent with the results of this study. The next step will be to develop a design scheme to improve a specific targeted peptide sequence in terms of affinity, biochemical properties, etc.

## 5. Conclusions

In this study, we applied the AfDesign binder hallucination protocol to PPI-targeting peptide design and attempted the simultaneous optimization of peptide sequence design and solubility control. Owing to the AfDesign flexible loss function design concept, we were able to design peptides likely to be PPI while controlling solubility by adding a solubility index to the loss. In addition, the evaluation of binding affinity by docking allowed us to design peptide sequences with higher binding affinity than the original peptides with PPI as the target protein. We suggest that further optimization of our approach could be applied to drug discovery through PPI-targeting peptide design.

## Author Contributions

Conceptualization, T.K. and M.O.; methodology, T.K. and M.O.; software, T.K.; validation, T.K.; formal analysis, T.K. and M.O.; investigation, T.K. and M.O.; writing—original draft preparation, T.K. and M.O.; writing—review and editing, T.K. and M.O.; visualization, T.K.; supervision, M.O.; project administration, M.O.; funding acquisition, M.O. All authors have read and agreed to the published version of the manuscript.

## Institutional Review Board Statement

Not applicable

## Informed Consent Statement

Not applicable

## Data Availability Statement

The implementation and experimental data are available on an opensource basis at https://github.com/ohuelab/Solubility_AfDesign (accessed on 13 May 2022).

## Acknowledgments

The computational experiments were performed using a TSUBAME 3.0 super-computer at the Tokyo Institute of Technology, for which we thank the institution.

