## Supplemental Materials for "Solubility-aware protein binding peptide design using AlphaFold"

#### 1. SUPPLEMENTARY FIGURES

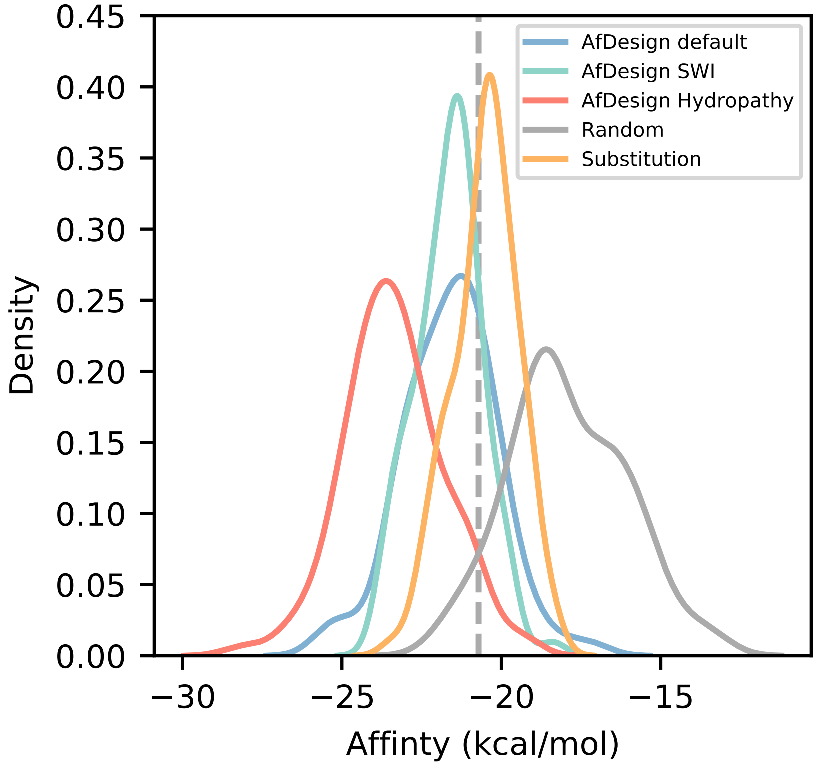


**Fig. S1.** Comparison of MDM2 binding affinities between random sequences and one residue substitution sequences of p53 peptides and sequences designed by AfDesign. The blue line shows the distribution of binding affinity of peptide sequences designed by default AfDesign, the green line shows the distribution of binding affinity of peptide sequences designed using the Solubility-Weighted Index as a solubility index, and the red line shows the distribution of binding affinity of peptide sequences designed using the Hydropathy Index as a solubility index. The orange line shows the distribution of binding affinity of peptide sequences, the orange line shows the distribution of binding affinity of one residue substitution sequences of p53 peptide sequences, the solid gray line shows the distribution of binding affinity of random sequences, and the dashed gray line shows the binding affinity value of the p53 peptide sequence.

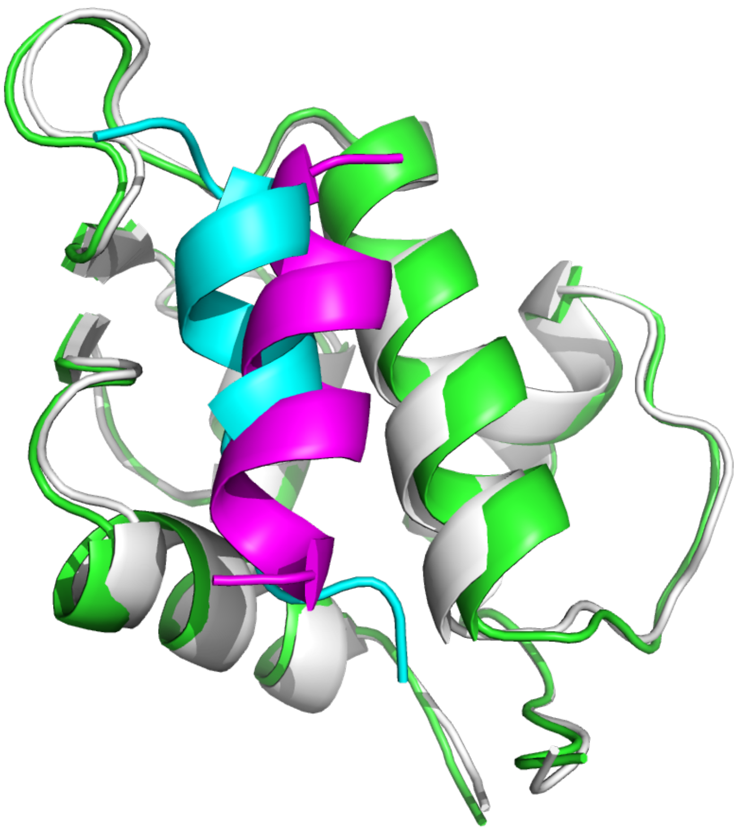

**Fig. S2.** Comparison of crystal structures and three-dimensional (3D) structures predicted by AfDesign. Green shows the crystal structure of MDM2, cyan shows the crystal structure of the p53 peptide, magenta shows the 3D structure of the sequence designed by AfDesign using the Solubility-Weighted Index as a solubility index (the sequence with the highest binding affinity and higher logS than that of the p53 peptide), and white shows the MDM2 structure as predicted by AfDesign.

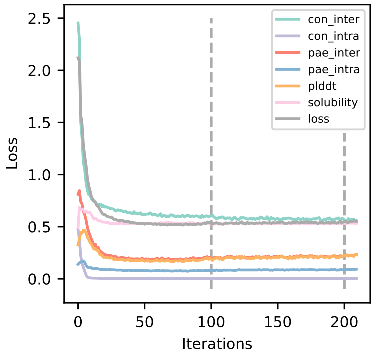


(**a**) Without solubility index

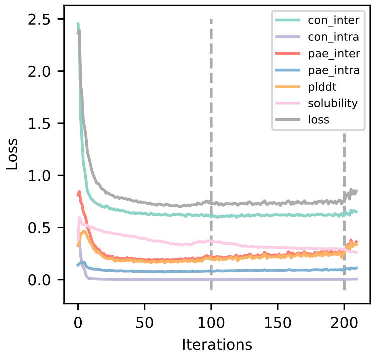


(**b**) Hydrophobicity index

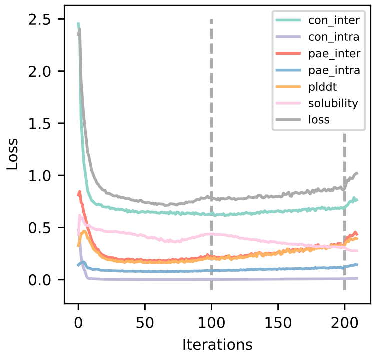


(**c**) Hydropathy index

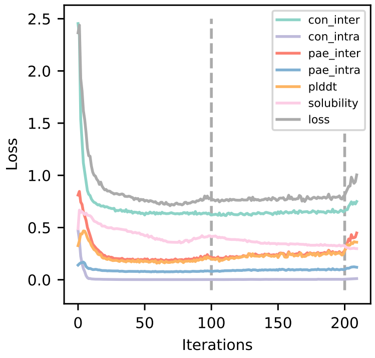


(**d**) SWI

**Fig. S3.** Comparison of each solubility index at the median of the AfDesign loss values. The green line shows the median loss value for each iteration of con_inter, the purple line shows the median loss value for each iteration of con_intra, the red line shows the median loss value for each iteration of pae_inter, the blue line shows the median loss value for each iteration of pae_intra, the orange line shows the median loss value for each iteration of plddt, the pink line shows the median loss value for each iteration of solubility, the solid gray line shows the median total loss weighted by the weighted average for each loss, and the dashed gray line shows the stage switching timing (101st, 201st iteration) in design_3stage() of the AfDesing binder hallucination protocol.

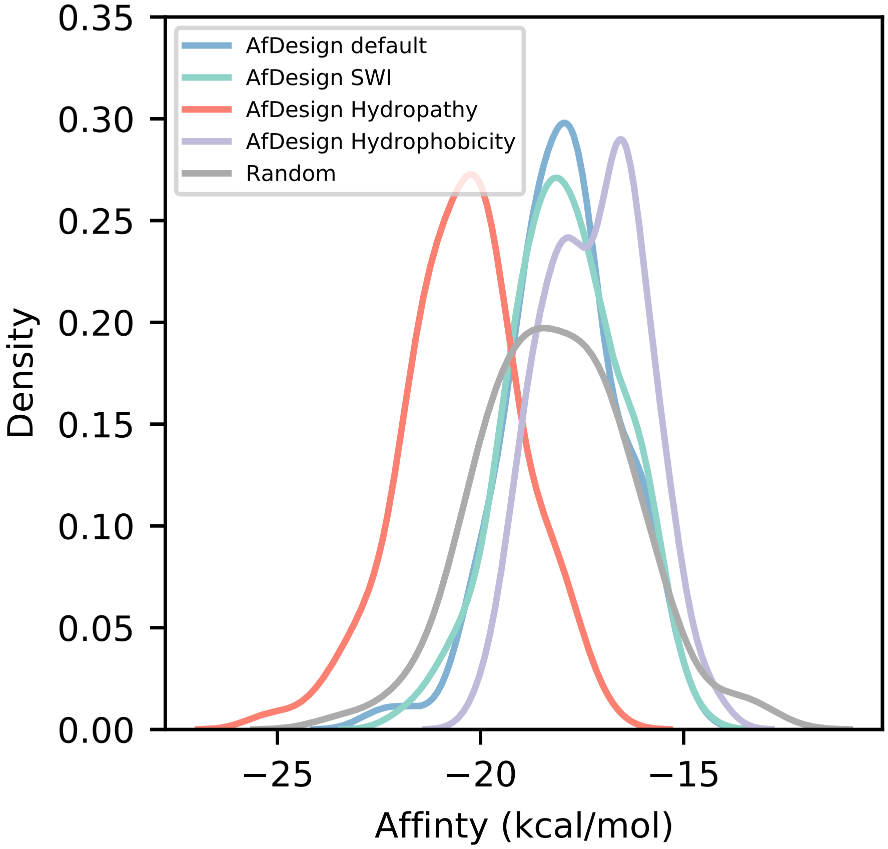


**Fig. S4.** Comparison of PD-1 binding affinity between random sequences and sequences designed in AfDesign with each solubility index. The blue line shows the distribution of binding affinity of peptide sequences designed by default AfDesign, the green line shows the distribution of binding affinity of peptide sequences designed using the Solubility-Weighted Index as a solubility index, the red line shows the distribution of binding affinity of peptide sequences designed using the Hydropathy Index as a solubility index, the purple line shows the distribution of binding affinity of peptide sequences designed using the Hydrophobicity Index as a solubility index, and the gray line shows the distribution of binding affinity of random sequences.

### 2. SUPPLEMENTARY TABLES

**Table S1.** Values of each solubility index. Values for the Hydrophobicity Index, Hydropathy Index, and Solubility-Weighted Index are referenced from each reference table. The leftmost column shows the one-letter code of amino acids.

| | Hydrophobicity Index | Hydropathy Index | Solubility-Weighted Index |
|---|---|---|---|
| A | 0.61 | 1.8 | 0.835647 |
| R | 0.60 | -4.5 | 0.771247 |
| N | 0.06 | -3.5 | 0.859743 |
| D | 0.46 | -3.5 | 0.907904 |
| C | 1.07 | 2.5 | 0.520809 |
| Q | 0.00 | -3.5 | 0.789435 |
| E | 0.47 | -3.5 | 0.987699 |
| G | 0.07 | -0.4 | 0.799717 |
| H | 0.61 | -3.2 | 0.894791 |
| I | 2.22 | 4.5 | 0.678412 |
| L | 1.53 | 3.8 | 0.655422 |
| K | 1.15 | -3.9 | 0.926710 |
| M | 1.18 | 1.9 | 0.629662 |
| F | 2.02 | 2.8 | 0.584979 |
| P | 1.95 | -1.6 | 0.823533 |
| S | 0.05 | -0.8 | 0.744091 |
| T | 0.05 | -0.7 | 0.809692 |
| W | 2.65 | -0.9 | 0.637468 |
| Y | 1.88 | -1.3 | 0.611280 |
| V | 1.32 | 4.2 | 0.735784 |

**Table S2.** Normalized solubility indices used as solubility loss for AfDesign. The three solubility indices were normalized to have a maximum of 1 and a minimum of 0, respectively. The Solubility-Weighted Index values were inverted (minus 1) after normalization. The leftmost column show the one-letter code of amino acids.

| | Hydrophobicity Index | Hydropathy Index | Solubility-Weighted Index |
|---|---|---|---|
| A | 0.230188679 | 0.700000000 | 0.325669858 |
| R | 0.226415094 | 0.000000000 | 0.463603847 |
| N | 0.022641509 | 0.111111111 | 0.274060271 |
| D | 0.173584906 | 0.111111111 | 0.170907494 |
| C | 0.403773585 | 0.777777778 | 1.000000000 |
| Q | 0.000000000 | 0.111111111 | 0.424648204 |
| E | 0.177358491 | 0.111111111 | 0.000000000 |
| G | 0.026415094 | 0.455555556 | 0.402625886 |
| H | 0.230188679 | 0.144444444 | 0.198993339 |
| I | 0.837735849 | 1.000000000 | 0.662440832 |
| L | 0.577358491 | 0.922222222 | 0.711681552 |
| K | 0.433962264 | 0.066666667 | 0.130628199 |
| M | 0.445283019 | 0.711111111 | 0.766855148 |
| F | 0.762264151 | 0.811111111 | 0.862558633 |
| P | 0.735849057 | 0.322222222 | 0.351616012 |
| S | 0.018867925 | 0.411111111 | 0.521767440 |
| T | 0.018867925 | 0.422222222 | 0.381261111 |
| W | 1.000000000 | 0.400000000 | 0.750136006 |
| Y | 0.709433962 | 0.355555556 | 0.806226306 |
| V | 0.498113208 | 0.966666667 | 0.539559639 |
